# Chronic hypertension and perfusion deficits conjointly affect disease outcome after tPA treatment in a rodent model of thromboembolic stroke

**DOI:** 10.1101/2024.03.20.585876

**Authors:** Bart AA Franx, Ivo ACW Tiebosch, Annette van der Toorn, Rick M. Dijkhuizen

**Author notes:** Correspondence: Bart A.A. Franx or Rick M. Dijkhuizen, Center for Image Sciences, University Medical Center Utrecht, Heidelberglaan 100, 3584 CX Utrecht, The Netherlands. or.

## Abstract

Futile recanalization hampers prognoses for ischemic stroke patients despite successful recanalization therapy. Allegedly, hypertension and reperfusion deficits contribute, but a better understanding is needed of how they interact and mediate disease outcome. We assessed data from spontaneously hypertensive and normotensive Wistar-Kyoto rats (male, n=6-7/group) that were subjected to two-hour embolic middle cerebral artery occlusion and thrombolysis in preclinical trials. Serial MRI allowed lesion monitoring and parcellation of regions-of-interest that represented infarcted (core) or recovered (perilesional) tissue. Imaging markers of hemodynamics and blood-brain barrier (BBB) status were related to tissue fate and neurological outcome. Despite comparable ischemic severity during occlusion between groups, hypertensive rats temporarily developed larger lesions after recanalization, with permanently aggravated vasogenic edema and BBB permeability. One day post-stroke, cerebral blood flow (CBF) was variably restored, but blood transit times were consistently prolonged in hypertensives. Compared to the core, perilesional CBF was normo-to- hyperperfused in both groups, yet this pattern reversed after seven days. Volumes of hypo- and hyperperfusion developed irrespective of strain, differentially associating with final infarct volume and behavioral outcome. Incomplete reperfusion and cerebral injury after thrombolysis were augmented in hypertensive rats. One day after thrombolysis, hypoperfusion associated with worsened outcomes, while regional hyperperfusion appeared beneficial or benign.

## Introduction

In acute ischemic stroke, poor outcome despite recanalization – or futile recanalization – is common and under (pre)clinical investigation to improve prognoses.^1^ Numerous hypotheses exist for futile recanalization,^1^ such as recanalization failure, re-occlusion, no-reflow, or more generally, post-recanalization perfusion deficits.^2^ These manifest as abnormal cerebral perfusion parameters after recanalization – i.e. hypo- or hyperperfusion – and are commonly reported in patients Accumulating clinical imaging data associate post-recanalization hypoperfusion with deterioration.^2–4^ Furthermore, hyperperfusion is reported regularly but prognostic value remains contested, possibly because it is an intricate function of time and spatial location.^2,4^

Translational imaging studies of reperfusion deficits indicate that hypertension may cause hypoperfusion upon recanalization.^2,5,6^ Hypertension is common in stroke populations^5^ and has been identified as a predictor of poor outcome after endovascular treatment.^7^ However, it remains unclear how hypertension affects post-recanalization hemodynamics and tissue outcome, particularly if there are temporal differences in reperfusion deficits between tissue areas bound for infarction or recovery. This was assessed in a clinically relevant model of embolic middle cerebral artery occlusion (eMCAO) and tissue plasminogen activator (tPA)-induced thrombolysis using serial multiparametric MRI. The objective was to identify features of reperfusion deficits that associated with worse post-ischemic tissue outcome in spontaneously hypertensive rats (SHRs) compared to normotensive rats (NRs).

## Methods

Data involved retrospective analyses of control groups used for blinded thrombolytic treatment studies^8^ (one unpublished). Imaging data used for these analyses is available at https://doi.org/10.5281/zenodo.10139231.

All animal procedures and experiments were approved by the Animal Experiments Committee of the University Medical Center Utrecht and Utrecht University and were performed in accordance with the guidelines of the European Communities Council Directive (protocol 2007.I.03.044/vervolg1 and 2008.I.3.031). They were performed in accordance with the Dutch Experiments on Animals Act and reported according to the ARRIVE guidelines.

### Animals

Normotensive male Wistar rats (300–380 g, aged 12–14 weeks) (Harlan, Venray, The Netherlands) or spontaneously hypertensive male Wistar-Kyoto (280–320 g, aged 12–14 weeks) (Charles River, Sulzfeld, Germany) were randomly allocated to their respective control group after an acclimatization period. Each rat represents one experimental unit. Rats were housed with a littermate under standard conditions, conforming to local legislation, with a light/dark cycle of 12/12 hours and free access to food and water.

### Data collection

Data presented in these retrospective analyses originate from two independent blinded thrombolytic treatment studies (one unpublished).^8^ Control groups of male normotensive rats (NRs) (n = 7) and male spontaneously hypertensive rats (SHRs) (n = 10) subjected to embolic stroke were selected for this analysis. Data sets were included based on two *a priori* criteria obtained from diffusion and perfusion MRI during the occlusion (baseline) phase of the experiment: 1) the ipsilateral tissue volume with an ADC two standard deviations below the mean contralateral value in anatomical areas fed by the MCA was larger than 30 µL, and 2) to exclude prematurely recanalized cases, mean transit time in the lesion core was more than 150% of the value in the contralateral homologous area (see *Regions-of-interest analyses* below for details). Only the second criterium led to the exclusion of four animals in the SHR group. Final group sizes were n = 7 (NR; control group) and n = 6 (SHR; experimental group). One animal in the NR group is lacks a third imaging session because, at the time of the preclinical trial, the lesion size was deemed too small to continue the experiment. Due to the retrospective nature of the study, randomization and blinding were not applicable.

### Animal handling

Rats were endotracheally intubated and mechanically ventilated with 2% isoflurane in an air- O_2_ mixture. Isoflurane anesthesia was maintained throughout microsurgery and MRI sessions. When anesthetized, body temperature was maintained at 37±0.5 °C by a feedback-controlled heating pad. Expired CO_2_ was monitored with a capnograph and kept within physiological range by manually changing the breathing rate and/or pressure of the rodent ventilator.

Prior to microsurgery, intramuscular injections of gentamicin (5 mg/kg) were given as an antibiotic. Animals received subcutaneous injections of 0.03 mg/kg buprenorphine (Temgesic®, Ricket & Colman, Kingston-Upon-Hill, UK) for pain relief and 2.5 ml glucose solution (2.5% in saline) to prevent dehydration. Post-operative care consisted of additional subcutaneous injections of 0.03 mg/kg buprenorphine and 2.5 ml glucose solution in saline (2.5%). Rats were weighed daily and excessive weight loss during the first three days was compensated with daily Ringer’s lactate (s.c., 0–10 ml, depending on lost weight).

### Experimental protocol

Right-sided unilateral embolic middle cerebral artery occlusion (MCAO) was induced as previously described.^8^ The right carotid artery was exposed by a ventral incision in the neck and a modified catheter was advanced into the internal carotid artery, until the tip was positioned just proximal to the MCA. An autologous blood clot (24 hours old) was slowly injected, followed by careful removal of the catheter. The incision was sutured and animals were prepared for MRI. After MCAO and the first MRI session, animals were treated with a tail vein infusion of 10 mg/kg tPA (Activase®, Genentech; concentrated to 3 mg/ml), of which 10% was delivered as a bolus, followed by continuous infusion of the remaining 90% over 30 minutes. According to their respective study protocol, NRs received an intravenous injection of phosphate-buffered saline directly after tPA treatment (i.v., once), while SHRs received daily subcutaneous injections of chemically inert vehicle substance (0.4% rat serum and 25 mmol/ml trehalose in PBS; pH 7) after stroke.

### Behavioral testing

Neurological status after stroke was assessed using the sensorimotor deficit score (SDS), an adapted battery of sensorimotor tests.^9^ The SDS was measured 1 day before stroke, and at days 1, 3, and 7 after stroke. Briefly, animals were scored consecutively on spontaneous exploratory mobility and gait disturbance, lateral resistance, forelimb placement guided by whisker stimulation, and forelimb grasping and -strength on a horizontal bar. The total SDS ranged from 0 (no deficit) to 22 (maximum deficit). Test items have been previously described.^10^ Researchers were blinded to treatment group allocation at the time of testing.

The pre-measurements were not further analyzed as no deficits were found. Day 3 was not analyzed because these data were not accompanied by MRI measurements.

### MRI

MRI was performed on a 4.7T horizontal bore MR system (Agilent, Palo Alto, CA, USA). A Helmholtz volume coil and inductively coupled surface coil (Ø 25 mm) were used for signal transmission and detection, respectively. Anesthetized rats were placed in a MR-compatible stereotactic holder and restrained with a headset and tooth bar, and mechanically ventilated. MRI was performed immediately after stroke induction (before tPA treatment), repeated at day 1 (Day 1) and day 7 (Day 7) after stroke. The imaging protocols for both groups, i.e. NRs and SHRs, were identical, with the exception of the MRI protocol for measurement of blood-brain barrier permeability (see below).

Diffusion-weighted 8-shot echo planar imaging (EPI; repetition time (TR) 3500 ms; echo time (TE) 38.5 ms; b-values 0 and 1428 s/mm^2^; 6 diffusion-weighted directions; field-of-view (FOV) of 32 × 32 mm^2^ with 1 mm slice thickness and a matrix size 128 × 128 × 19) was performed. Sequential 2D spin-echo T_2_-weighted images (TR/[TE] 3600/[12–144] ms, 12 equidistant echoes), and T_2_*-weighted images (TR/[TE] 1400/[7–70] ms; 10 equidistant echoes), both with an FOV of 32 × 32 mm^2^ with 1 mm slice thickness and a matrix size 256 × 128 × 19 were acquired for reconstruction of quantitative T_2_- and T_2_*-maps. To measure cerebral perfusion parameters, dynamic susceptibility contrast-enhanced (DSC) MRI, using a 2D gradient echo EPI (TR/TE 330/25 ms; 400 dynamics; matrix size = 64 × 64 × 5) was combined with a tail vein bolus injection of 0.35 mmol/kg gadobutrol (Gadovist®, Schering, The Netherlands) after 30 seconds. Lastly, magnetic resonance angiography was performed using a flow-compensated 3D time-of-flight sequence (gradient echo; TR/TE 15/2.66 ms; FOV = 32 × 32 × 32 mm^3^; matrix size = 128 × 128 × 128).

To assess blood-brain barrier leakage in SHRs, T_1_-weighted images were acquired (2D gradient echo; TR/TE 160/4 ms; flip angle 90°; FOV of 32 × 32 mm^2^ with 1 mm slice thickness; data matrix 256 × 128 × 19) every 2.73 min up to 35 min after gadobutrol injection. For NRs, a 2D gradient echo EPI Look-Locker sequence was employed (TR/TE 3000/4 ms; 28 equidistant inversion times per slice ranging from 9 to 1809 ms; flip angle 10°, FOV of 32 × 32 mm^2^ with 2 mm slice thickness; matrix size = 128 × 64 × 5).

### Image processing

Mean apparent diffusion coefficient (ADC) maps were obtained after fitting the full tensor of the diffusion matrix. Quantitative T_1_-, T_2_- and T_2_*-maps were calculated by derivative-based separable least-squares fitting of complex-valued data. For SHRs, quantitative T_1_- mapping was not performed, but T_1_-weighted images were collected. As described earlier,^11^ these images were used to calculate T_1_-maps using the relationship

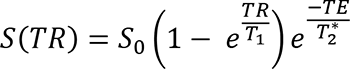

with S_0_ as the estimated proton density obtained from T_2_-maps. R_1_ maps (reciprocal of T_1_) were calculated for both groups and used to estimate the blood-to-brain transfer constant (K_i_) and distribution space of intravascular Gd-shifted protons (V_p_) using the Patlak graphical analysis approach.^12^ The plasma concentration was estimated from the sagittal sinus (twenty voxels). For NRs, T_1_ was measured from magnetization recovery in the Look-Locker experiment.

There was a small but systemic difference in R_1_ between SHRs and NRs, possibly due to T_1_ being estimated differently (see above). The R_1_ offset propagates into the K_i_ calculations, and as such, absolute K_i_ values could not be compared directly. Relative K_i_ values were therefore calculated by expressing the mean K_i_ from an ipsilesional region as a percentage of the mean K_i_ from the contralesional homologue (see *Region-of-interest analysis* below).

Cerebral blood flow (CBF) maps were calculated by circular deconvolution of tissue concentration curves using a contralesional arterial reference curve.^13^ Cerebral blood volume (CBV) was calculated by numeric integration of the tissue concentration curve. Mean transit time (MTT) was the quotient of CBV and CBF.

Using the ANTs software package,^14^ MRI images were corrected for B_1_ inhomogeneity and coregistered. Brain extraction was performed with FSL’s Brain Extraction Tool.^15^ All T_2_-weighted images with the least T_2_-weighting (i.e., shortest TE) served as anatomical scans and were co-registered ‘inter-session’ to a study-specific template space, while also being used ‘intra-session’ as a reference for co-registration of parametric maps (i.e., ADC-, T_2_-, perfusion, and K_i_-maps) and lesion masks. By doing so, all data from both groups could be aligned and analyzed in a common space (see below). Non-linear registration was applied to correct space-occupying edema (characterized by midline shift) or deformations caused by gradient echo sequences.

### Region-of-interest analyses

To track lesion evolution, semi-automatic delineation of cytotoxic edema on ADC-maps during occlusion (*acute ischemia*) and vasogenic edema on T_2_-maps at Day 1 and Day 7 (*Day 1 infarct* and *Day 7 infarct*) were performed as described previously.^16^ In brief, each anatomical scan from each time point was coregistered to a normalized rat brain that contained a parcellated image of the Paxinos and Watson brain atlas^17^ which could then be backprojected to the ADC-map (occlusion) and T_2_-maps (Day 1 and Day 7). ADC and T_2_ values were extracted from the contralateral MCA territory. Their mean (*x̄*) and standard deviation (*sd*) were calculated, and acute ischemic tissue was defined by an ADC at least 2 standard deviations lower than the mean contralateral ADC (< x̄ − 2*sd*), while for T_2_-maps, voxels were considered vasogenic edematous if the T_2_ exceeded the contralateral mean by more than 2 standard deviations (> x̄ + 2*sd*). Lesion masks were manually noise-corrected.

Then, in template space, the following ROIs were derived for each subject using basic set operations:

1. irreversible acute ischemic injury during occlusion that proceeded to infarction at Day 7, termed the *lesion core,* i.e. voxels that were *acute ischemic area* ∩ *Day 7 infarct*
2. reversible perilesional injury, tissue that exhibited prolonged T_2_ at Day 1 but did not proceed to infarction at Day 7, termed *perilesional tissue,* i.e., voxels that were *Day 1 lesion* ∪ *Day 7 infarct* Contralateral homologues were obtained by aligning mirrored copies of these ROIs to a mirrored copy of the common template reference image.

In common template space, mean ADC- and T_2_-values were extracted from the ipsilateral ROIs described above. Regarding perfusion maps, mean values from ipsilateral ROIs were expressed as a percentage of their contralateral counterparts, producing a relative regional perfusion index (rrCBF, rrCBV and rrMTT). This process eliminates inadvertent intra- and inter-subject variability stemming from bolus injection efficiency. The same normalization process was applied to values from K_i_-maps, since the distinct R_1_-estimation methods applied to the normotensive and hypertensive groups produced a systemic difference in K_i_ maps between groups (see *Image processing* above).

Lesion sizes were calculated in template space. The total number of voxels that a lesion mask was comprised of, for each experimental time point, was multiplied by the voxel dimensions (spatial resolution). Each individual lesion mask was normalized by alignment to the common study template image, therefore obviating the need for hemispheric lesion volume calculations. Hence, lesion volumes in µL were obtained.

Volumes of post-recanalization perfusion deficits (reperfusion deficit), i.e., hypo- and hyperperfusion, were objectively quantified as described in earlier work.^3^ The vasogenic edematous lesion on Day 1 exhibited much variability in reperfusion deficit and was therefore of main interest. Briefly, we operationalized hypo- and hyperperfusion for each cerebral perfusion parameter by binning each voxel in the ipsilateral vasogenic edematous lesion based on a comparison to a fair control, here the mean obtained from the contralesional homologous mask. It was recently shown that a perfusion asymmetry of ±15% can be clinically significant.^3^ Thus, if a voxel in the vasogenic edematous lesion was >115% of the contralateral mean, it was identified as hyperperfused. Analogously, if it was <85% of the contralateral mean, it was hypoperfused. These definitions were used to identify reperfusion deficit on CBF and CBV maps, but for MTT maps the definitions were reversed: short (<85%) and long (>115%) MTT values signified hyper- and hypoperfusion, respectively. Voxels within these boundaries were identified as normoperfused. Hypo- and hyperperfused voxels were counted, and their volume was expressed as a percentage of the subject’s vasogenic edematous tissue visible on the perfusion scan, to estimate the fraction of reperfusion deficit that occupied it. This normalization step was necessary, because the extent of vasogenic edema varied considerably between subjects and also partially existed outside of the field-of-view of the DSC-MRI experiment, thus rendering absolute volumes of reperfusion deficit meaningless for further analysis.

### Statistical analyses

Assumptions for each parametric test, such as the supposed distribution of the response variable (i.e., normal or beta), homogeneity of variance and normality of residuals, were checked but no violations were encountered. Degrees-of-freedom were approximated by the Kenward-Roger method in mixed models. In linear mixed-models, Subject (i.e. rat) was always set as random effect. For multiplicity corrections in post-hoc testing, Bonferroni was applied at all times.

To analyze differences in lesion size over time, linear mixed-model analysis was performed for the size of the lesion measured during occlusion (cytotoxic edema; ADC-map), at Day 1 (vasogenic edema; T_2_-map) and at Day 7 (*idem*). Time (occlusion, Day 1, Day 7) and Strain (SHR vs. NR) and their interactions were considered fixed effects.

Imaging markers of cerebral injury in the lesion core ROI were analyzed by fitting linear mixed-models using mean ADC (cytotoxic edema), T_2_ (vasogenic edema) and the blood-to-brain indicator transfer constant (K_i_; blood-brain barrier permeability) as response variables. Time (occlusion, Day 1, Day 7) and Strain (SHR vs. NR) and their interaction were considered fixed effects.

To analyze differences in perfusion magnitude in areas of certain tissue fate over time, linear mixed-model analysis was performed for each relative regional perfusion index. rrCBF, rrCBV and rrMTT were compared by Strain (SHR vs. NR), Time (occlusion vs. Day 1 vs. Day 7) and region-of-interest (ROI; lesion core vs. perilesional tissue).

To analyze the association between fractions of reperfusion deficit (hypo- and hyperperfusion) occupying the vasogenic lesion at Day 1 on the infarct size, ordinary least-squares fits were performed using the vasogenic lesion size at Day 7 as response variable. Strain (SHR vs. NR) was a cofactor. Initial size of the volume of cytotoxic edema (acute ischemia), as measured on the first ADC map, was a control variable. Principally the same strategy was applied to test the effect of fractional reperfusion deficit on neurological outcome after seven days, except that a generalized linear model was employed, where a Poisson distribution of the dependent variable was assumed. Either the fraction of hypo- or hyperperfusion was included as a covariate to estimate their contribution to stroke outcome: models were fitted separately due to the strong negative correlation between fractions of reperfusion deficit for a range of thresholds (Supplementary Figure I(a)). For a ±15% threshold, the correlation between hypo- and hyperperfusion fractions in the vasogenic lesion was near-perfect. Including both parameters would produce unreliable models with unacceptable variation inflation factors and unstable coefficients. Instead, one may consider that with multicollinearity of this degree, either hypo- or hyperperfusion is redundant in the regression equation.

Statistical analyses were performed in R 4.0.2 using the *tidyverse,*^18^ *lme4,*^19^ *emmeans,*^20^ *car,*^21^ *ggtern,*^22^ and *modelsummary*^23^ packages.

## Results

One day after eMCAO, vasogenic edematous lesions (i.e., with T_2_ prolongation) were larger in SHRs compared to NRs (*p* = .03; Figure 1(a)). The mean SDS was also higher at this time but not significantly so (*p* = .07; Figure 1(b)).

**Figure 1.**
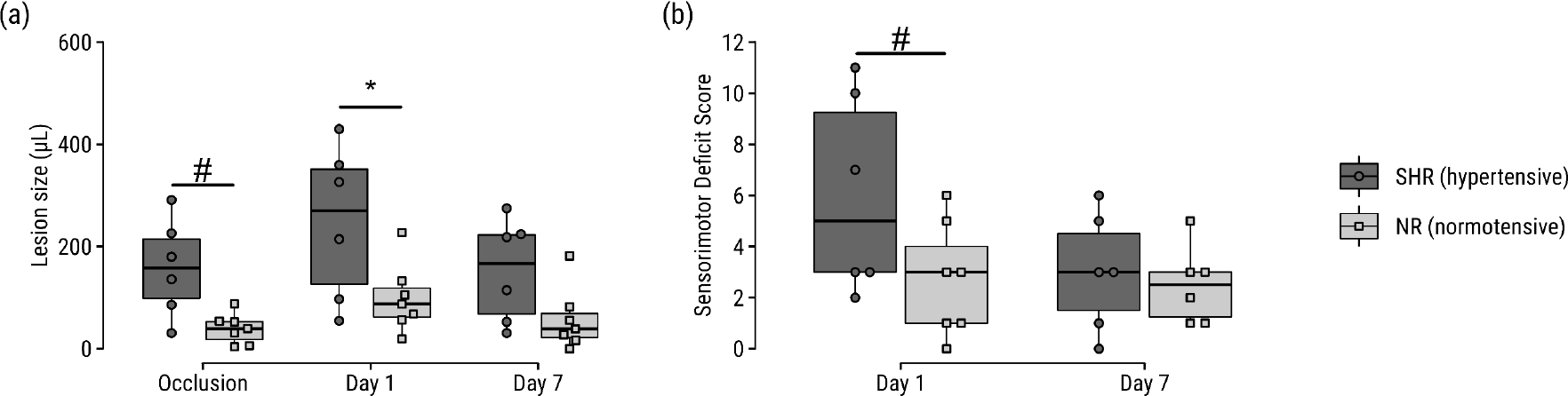
Increased lesion size and sensorimotor deficit score in SHRs compared to NRs. (a) Lesion volumes before and after tPA-treated eMCAO, based on diffusion- and T_2_ MRI of cytotoxic and vasogenic edema, respectively. (b) Sensorimotor deficit scores at Days 1 and 7 after tPA-treated eMCAO. \**p*<0.05. ^#^*p*<0.10

Serial multiparametric MRI enabled close monitoring from ischemia until infarction (Figure 2(a)). Regions-of-interest were parcellated (Figure 2(b)) into *core* (ischemic tissue still injured at follow-up) and *perilesional* tissue (non-ischemic yet injured at Day 1 but ultimately recovered at Day 7). In the core, no differences were found in level of ADC reduction during occlusion between groups (Figure 2(c)). T_2_ prolongation in the core was increased in SHRs on Day 1 (*p* = .03) and Day 7 (*p* < .0001). The blood-to-brain indicator transfer constant was elevated in SHRs at Day 1 (*p* = .0006) and Day 7 (*p* = .02).

**Figure 2.**
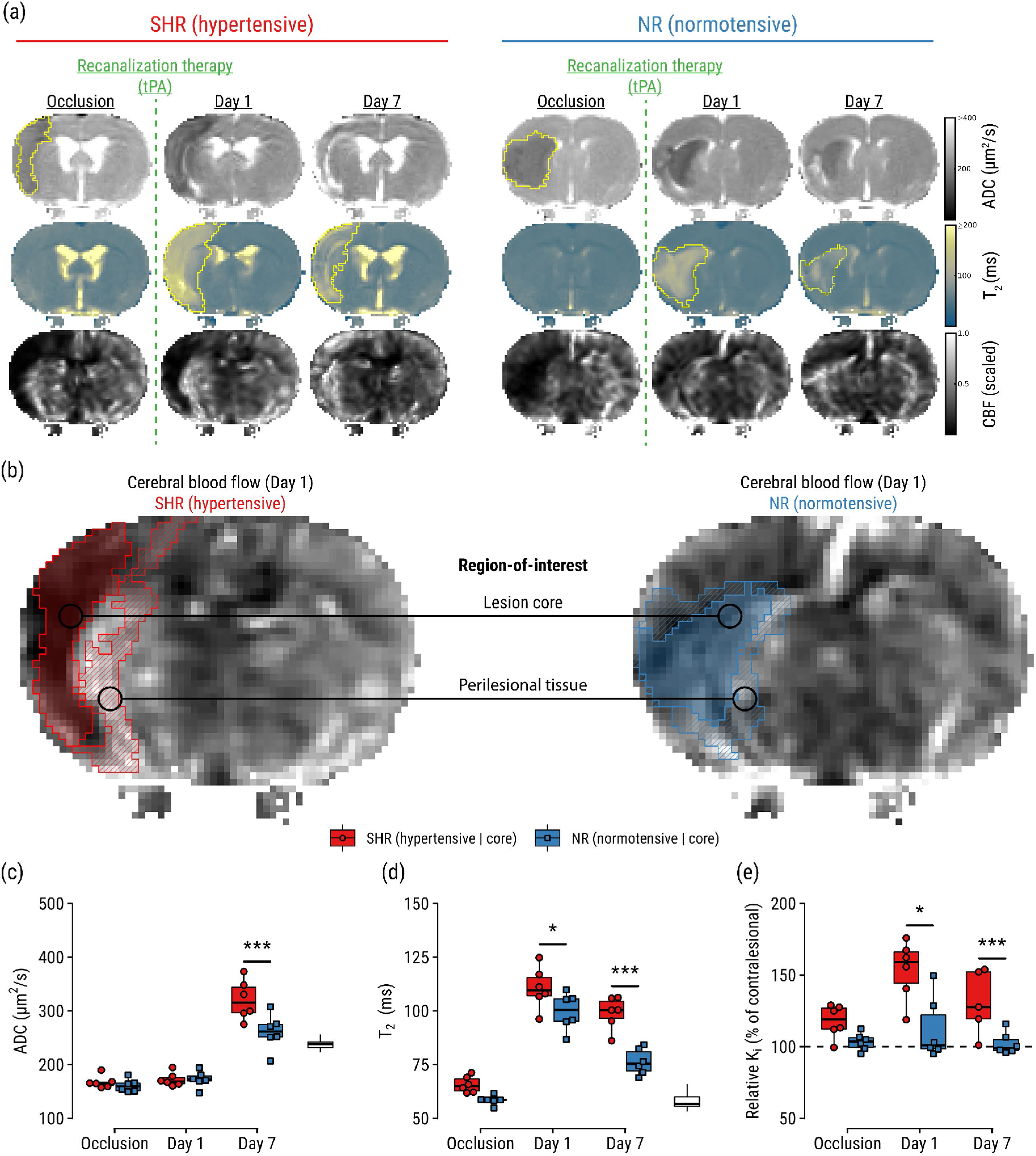
Region-of-interest analyses guided by imaging markers of stroke outcome, which are aggravated in SHRs. (a) Representative maps of apparent diffusion coefficient (ADC), T_2_, and cerebral blood flow (CBF) of a hypertensive and normotensive control subject. Lesions were semi-automatically segmented on ADC-maps during occlusion and on T_2_-maps at Day 1 and Day 7. (b) Regions-of-interest: lesion core defined by ADC reduction (cytotoxic edema) during occlusion and T_2_ prolongation (vasogenic edema) on Day 7; and perilesional tissue defined by normal ADC during occlusion, T_2_ prolongation on Day 1, and normal T_2_ on Day 7. Regions-of-interest are overlaid on CBF-maps showing post-recanalization perfusion status at Day 1. (c) ADC, T_2_ and relative blood-to-brain transfer constant (*K_i_*) in the lesion core. White boxplots indicate contralateral values for reference. \**p*<0.05, \*\*\**p*<0.001.

The eMCAO model exhibited variable tPA-induced reperfusion levels on Day 1 (Figure 3). A significant interaction was detected for rrCBF over time: while rrCBF was lower in the core than in perilesional tissue at Day 1, the opposite was observed at Day 7 for both SHRs (*p* = .0004) and NRs (*p* = .02) (Figure 3(a), top). Additionally, an interaction effect indicated that the difference in CBF between core and perilesional tissue seen at Day 1 switched sign on Day 7 for both groups, suggesting that perilesional areas were higher perfused than core areas on Day 1, but this pattern reversed on Day 7 (Supplementary Table III). Similar patterns were found for rrCBV, with stronger higher perfusion in the core compared to perilesional tissue in SHRs at Day 7 (*p* = .008; Figure 3(a), middle). Lastly, rrMTT at Day 1 was prolonged in the core of SHRs compared to NRs (*p* = .02; Figure 3(a), bottom), and the difference in rrMTT between lesion cores and perilesional tissue was unequal between both groups (*p* = .03). See Supplementary Tables I-III for omnibus and relevant post-hoc tests.

**Figure 3.**
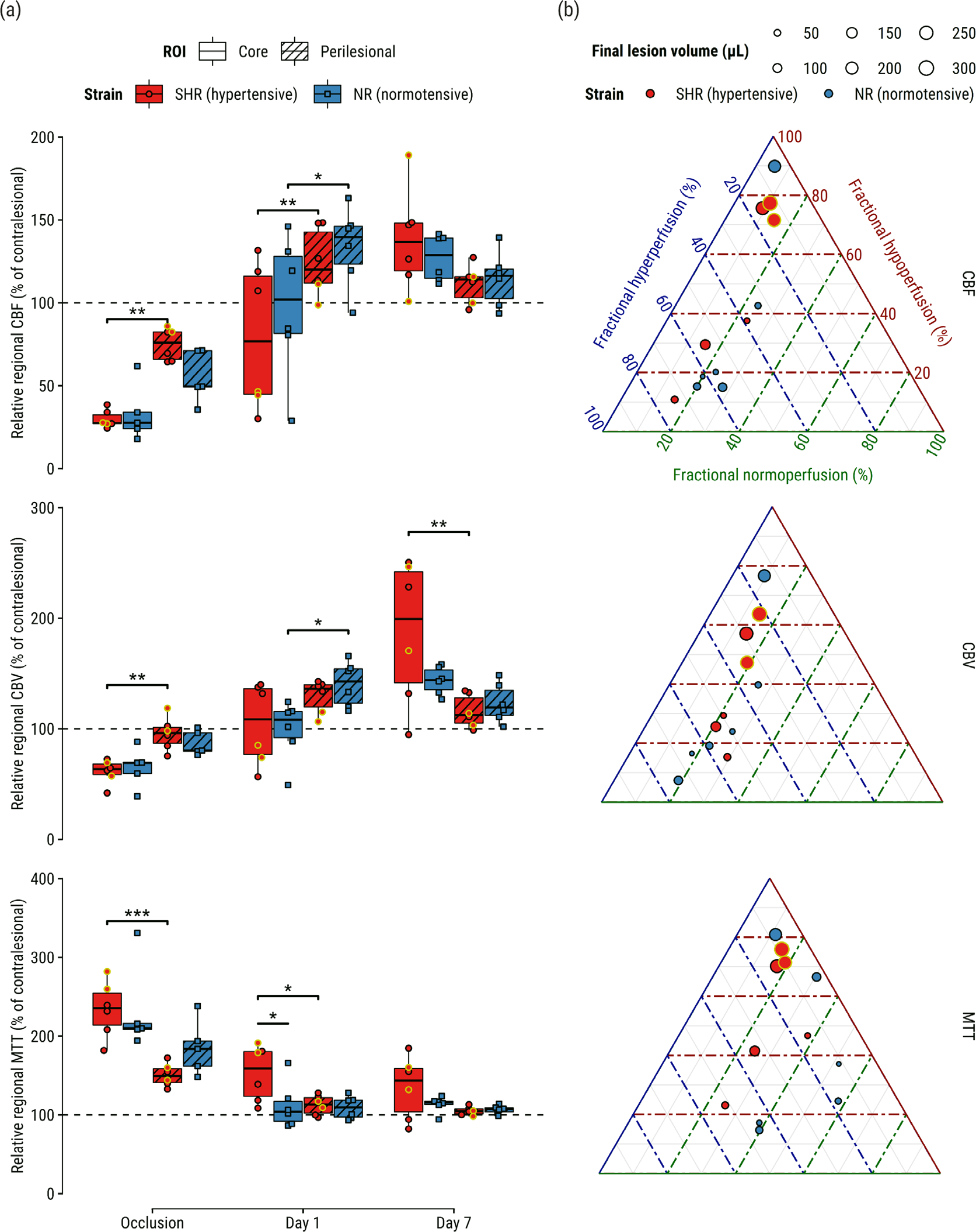
Perfusion abnormalities after tPA-treatment associate with hypertensive phenotype, tissue fate-derived regions-of-interest, and infarct size. (a) Relative regional cerebral blood flow (CBF) (top), cerebral blood volume (CBV) (middle), and mean transit time (MTT) (bottom). (b) Ternary plots visualizing fractions of hypo- hyper- and normoperfused tissue in vasogenic edematous lesion at Day 1 (center T_2_-maps in Figure 2A) for CBF (top), CBV (middle), and MTT (bottom). Each dot represents a subject; its size indicates infarct size at Day 7. Yellow-outlined dots represent subjects with impaired reflow at Day 1; Supporting angiograms (Supplementary Figure II) show recanalization of the previously occluded artery. \**p*<0.05, \*\**p*<0.01, \*\*\**p*<0.001.

Reperfusion deficit fractions in the T_2_ lesion on Day 1 showed near-perfect negative correlation (Supplementary Figure I(a)), indicating that lesions at Day 1 were primarily hypo- or hyperperfused. There were no differences in reperfusion deficit fractions between groups (Supplementary Figure I(b)). Higher fractions of hypoperfusion, expressed by CBF, CBV or MTT, were associated with larger infarcts (*p* < .01, *p <* .05 and *p* < .001, respectively). Conversely, higher fractions of CBF hyperperfusion were associated with smaller final lesion volumes (*p* < .05), but this relationship was non-significant for the other perfusion indices (Table I). Regarding neurological outcome, higher fractional hypoperfusion in terms of MTT in the vasogenic lesion at Day 1 associated with an increased SDS at Day 7 (Supplementary Table IV).

**Table I:**
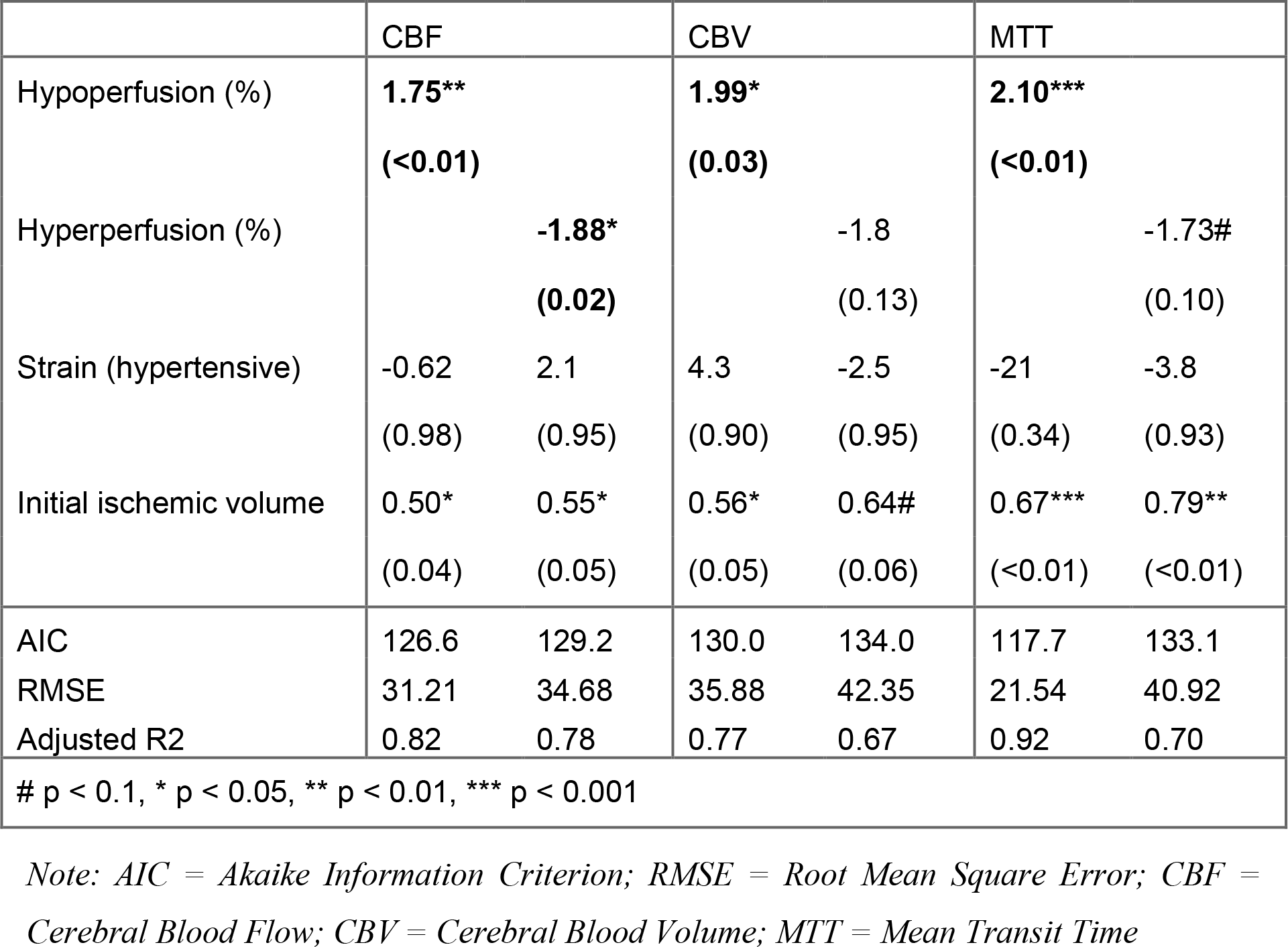
fractions of hypo- and hyperperfusion regressed against final infarct size.

|  | CBF |  | CBV |  | MTT |  |
| --- | --- | --- | --- | --- | --- | --- |
| Hypoperfusion (%) | **1.75\*\*** |  | **1.99\*** |  | **2.10\*\*\*** |  |
|  | **(<0.01)** |  | **(0.03)** |  | **(<0.01)** |  |
| Hyperperfusion (%) |  | **-1.88\*** |  | -1.8 |  | -1.73# |
|  |  | **(0.02)** |  | (0.13) |  | (0.10) |
| Strain (hypertensive) | -0.62 | 2.1 | 4.3 | -2.5 | -21 | -3.8 |
|  | (0.98) | (0.95) | (0.90) | (0.95) | (0.34) | (0.93) |
| Initial ischemic volume | 0.50* | 0.55* | 0.56* | 0.64# | 0.67*** | 0.79** |
|  | (0.04) | (0.05) | (0.05) | (0.06) | (<0.01) | (<0.01) |
| AIC | 126.6 | 129.2 | 130.0 | 134.0 | 117.7 | 133.1 |
| RMSE | 31.21 | 34.68 | 35.88 | 42.35 | 21.54 | 40.92 |
| Adjusted R2 | 0.82 | 0.78 | 0.77 | 0.67 | 0.92 | 0.70 |
| # p < 0.1, * p < 0.05, ** p < 0.01, *** p < 0.001 |  |  |  |  |  |  |

*Note: AIC = Akaike Information Criterion; RMSE = Root Mean Square Error; CBF = Cerebral Blood Flow; CBV = Cerebral Blood Volume; MTT = Mean Transit Time*

The relationship between fractional reperfusion deficit in the lesion on Day 1 and final infarct volume at Day 7 is illustrated in ternary plots for CBF, CBV and MTT (Figure 3(b)). Interestingly, areas of hyperperfusion seen on CBF and CBV maps tended to cancel out when expressed as MTT (Figure 3(a-b)).

## Discussion

Here, it was shown that despite comparable ischemic severity during embolic occlusion, SHRs develop larger lesions one day after treatment with tPA, with lasting aggravated edema and BBB permeability. Through exhaustive assessment of cerebral hemodynamics, dissected into three key variables – CBF, CBV and MTT – it was found that blood transit times were notably prolonged in lesion cores of the SHRs. Further, distinct hemodynamic patterns were found in tissues of foregone destruction or destined for recovery. Increased perfusion was clearly present in perilesional areas on Day 1, but these areas later exhibited recovery at Day 7 (T_2_-normalization). Accordingly, perilesional areas have previously been associated with resolution of edema^24^ and glial proliferation.^25^ Lastly, it was found that hypoperfusion at Day 1 associated with lesion growth and worsened behavioral deficits, while hyperperfusion seemed harmless.

Importantly, our findings support the notion that early hyperperfusion after recanalization could be benign, while late hyperperfusion may be a sequel of ischemic injury.^2^ Furthermore, in our rat model, early hyperperfusion was mostly present *around* the lesion, while perfusion only later became excessive in the lesion core, which (by definition) had not recovered on day 7. The recognition that peri-lesional hyperperfusion is associated with good outcomes originates from clinical literature.^2,26^ While hyperperfusion is well-described in animal models of stroke at various moments post-recanalization, translational imaging studies have investigated or framed it exclusively in light of injurious processes after ischemia.^2^ Little is still known about the processes that drive hyperperfusion early after recanalization in peri-lesional areas. Interestingly however, magnetic resonance spectroscopy in tPA-treated stroke patients who exhibited hyperperfused peri-lesional tissue one day after stroke – which associated with better 3-month outcome – revealed a metabolic signature that suggested increased cellular activity.^27^ In patient studies, it remains unclear if acutely hyperperfused tissue proceeds to infarction or recovery because subacute or chronic time points are typically not included. The present data adds to the limited body of knowledge by showing 1) hyperperfused peri-lesional tissue exhibits signs of recovery after the acute phase of thrombo-embolic stroke in normo- and hypertensive rats, and 2) peri-lesional hyperperfusion can be replicated for investigation in an experimental stroke model, such that tissue recovery processes can be better understood.

Our results also further add to the growing body of literature describing far-reaching effects of vascular comorbidity on stroke outcome.^5^ The translational axis is often decried due to numerous failures of experimental treatments in the clinic for a plethora of reasons (e.g., including only young healthy males in preclinical trials). Simultaneously, certain candidate cerebroprotective strategies already fail in preclinical studies incorporating comorbidity.^28,29^ The association between vascular comorbidity and impaired reperfusion provides a plausible explanation, not only for why recanalization therapy fails, but it would also hamper candidate drugs in reaching the lesioned parenchyma after recanalization.

Therefore, it is conceivable that cerebroprotective strategies require effective reperfusion in both stroke model and patient. The association between hypertension and persistent hypoperfusion deficiency should be verified in patient populations, while preclinically, treatment strategies that respond to comorbidities that affect the neurovascular unit are highly anticipated.^30^

Several pathomechanisms are implicated that can mediate impaired perfusion, brain edema and BBB permeability after stroke in SHRs. The hypertensive phenotype may exacerbate brain edema and BBB breakdown after stroke as a result of dysfunctional tight-junction protein expression^31^ and post-stroke inflammatory responses.^32^ Furthermore, SHRs have altered vascular structure and function compared to their normotensive counterparts. Multiparametric MRI has shown that the cerebrovascular reserve of SHRs is inherently weak, which may have contributed to the observed effects.^33^ In line with this, pial collateral vessels of SHRs have shown reduced diameters and increased vascular tone before, during and hours after filament-induced ischemia.^34^ This effect may persist long after the initial time window of recanalization to continuously hamper reperfusion as seen in the present analysis.

Certain limitations confine these results to being mostly hypothesis-generating, particularly due to the retrospective nature of the dataset, which also precluded appending additional experimental groups, such as female SHRs. Also, while chronic high blood pressures are consistently described in SHRs, this was unfortunately not verified at the time of data collection. Retrospective data collection is prone to certain biases, but it should be noted that investigator’s bias was mitigated by automating the analysis pipelines presented here. Importantly, with the re-use of historical control groups, this research respects the 3Rs in animal experimentation.

In conclusion, longitudinal MRI imaging of transient eMCAO in spontaneous hypertensive and normotensive rat strains revealed that congenital hypertension associates with worsened disease outcomes in terms of acute lesion size, edema, and BBB permeability. Incomplete (delayed) reperfusion was present in the lesion cores of hypertensive rats one day after thrombolysis. At the same time, regardless of strain, hyperperfusion occurred in peri- lesional areas that would ultimately recover. In line with this, it was shown that fractional hyperperfusion in the day old vasogenic lesion was harmless, while higher fractional hypoperfusion associated with lesion growth and neurological impairment. It will be of particular interest to measure similar outcomes in other models of acquired hypertension; preferably in conjunction with antihypertensive treatment, which may ameliorate the detrimental effects of hypertension on stroke outcome. Future studies that tackle these matters can make substantial contributions towards preventative medicine and advances in recanalization therapies.

## Author contribution statement

R.M.D., and B.A.A.F. designed the study. I.A.C.W.T. performed microsurgeries and acquired MRI data. A.vd.T. designed MRI sequences and contributed to acquisition. All authors contributed to data analyses and interpretation. B.A.A.F. and R.M.D. drafted the manuscript. All authors were involved in final interpretation and revising of the manuscript.

## Acknowledgements

The authors thank Gerard van Vliet for excellent technical support.

## Disclosures

The authors report no conflicts.

## Supplementary material

Supplementary material can be found at …

## Funding

The CONTRAST consortium acknowledges the support from the Netherlands Cardiovascular Research Initiative, an initiative of the Dutch Heart Foundation (CVON2015- 01: CONTRAST), and from the Brain Foundation Netherlands (HA2015.01.06). The collaboration project is additionally financed by the Ministry of Economic Affairs by means of the PPP Allowance made available by the Top Sector Life Sciences & Health to stimulate public–private partnerships (LSHM17016 and LSHM17020). This work was funded in part through unrestricted funding by Stryker, Medtronic, and Cerenovus.

## Notes

https://doi.org/10.5281/zenodo.10139231

### Competing Interest Statement

The authors have declared no competing interest.

