## Supplementary for "Chronic hypertension and perfusion deficits conjointly affect disease outcome after tPA treatment in a rodent model of thromboembolic stroke"

### Supplementary material

Franx, et al.

2024-02-21

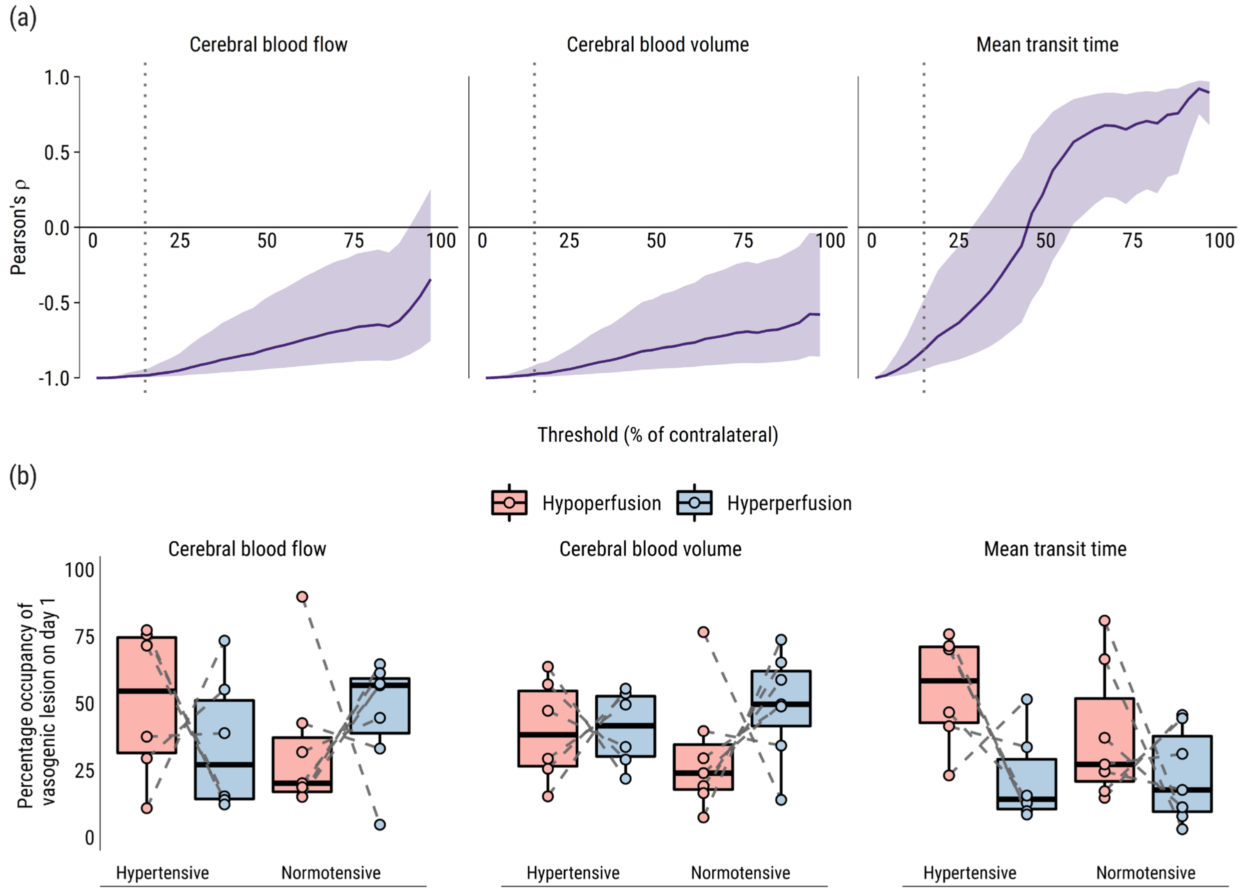


**Supplementary Figure 1.** The correlation between reperfusion deficiencies in the vasogenic lesion (day 1) and their definition. (a) Pearson's correlation coefficients between hypo- and hyperperfusion fractions, plotted as a function of threshold values. Shaded areas represent ±95% confidence interval. Dotted grey lines denote 15% threshold used for this analysis. (b) Percentages of hypo- and hyperperfused areas present within the vasogenic edematous lesion on day 1 (Figure 2(a)) using a 15% threshold. Dashed interconnected lines represent within-subject measurements, illustrating the origin of the strong negative correlation between hypo- and hyperperfusion measured in this sample.

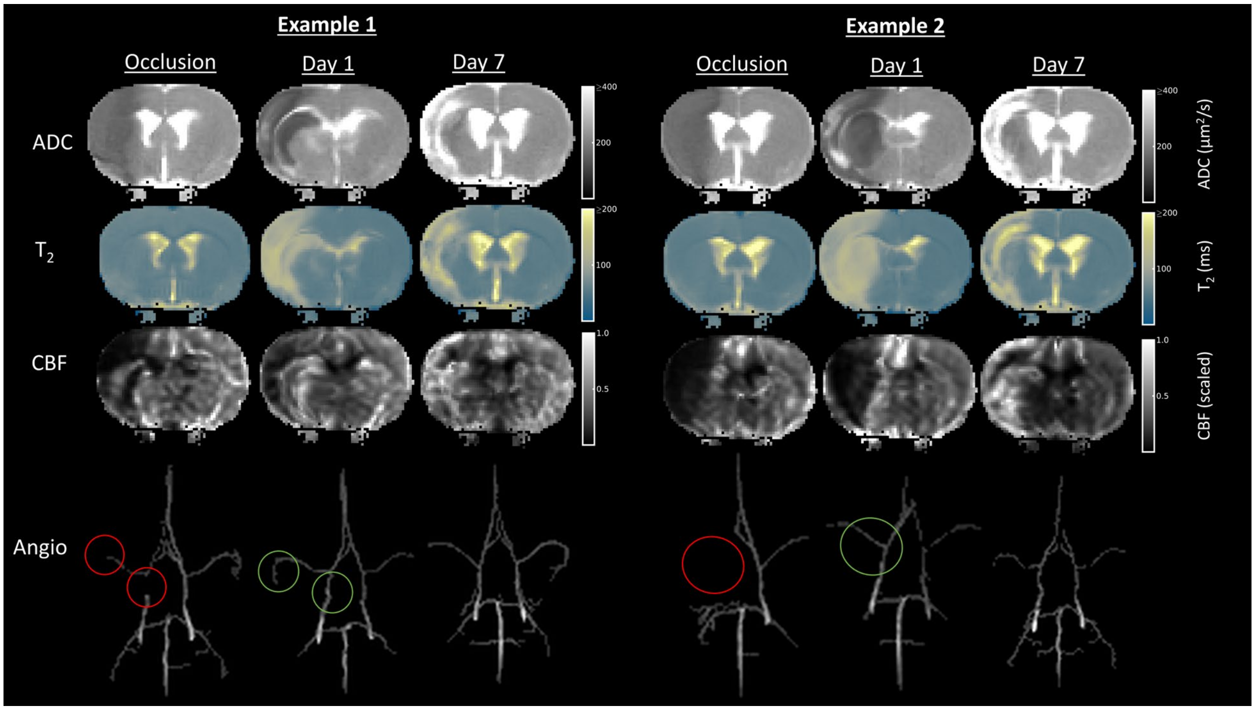


**Supplementary Figure 2.** Serial coronal brain maps of ADC, T2 and CBF, and MR angiograms from two SHRs with recanalized feeding arteries but impaired reperfusion on day 1.

**Supplementary Table I - RM-ANOVA – summary of linear mixed model perfusion analysis**

CBF

| Term | F | Df | Residual df | *p* |
|---|---|---|---|---|
| Strain | 0 | 1 | 9.95 | > 0.99 |
| ***Time*** | ***77.05*** | ***2*** | ***48.51*** | ***< 0.001*** |
| ***ROI*** | ***11.93*** | ***1*** | ***48.02*** | ***0.001*** |
| Strain:Time | 2.09 | 2 | 48.59 | 0.13 |
| Strain:ROI | 0.28 | 1 | 48.02 | 0.6 |
| ***Time:ROI*** | ***14.02*** | ***2*** | ***48.02*** | ***< 0.001*** |
| Strain:Time:ROI | 1.07 | 2 | 48.02 | 0.35 |

CBV

| Term | F | Df | Residual df | *p* |
|---|---|---|---|---|
| Strain | 0.18 | 1 | 9.93 | 0.68 |
| ***Time*** | ***45.56*** | ***2*** | ***48.68*** | ***< 0.001*** |
| ***ROI*** | ***6.19*** | ***1*** | ***48.04*** | ***0.016*** |
| Strain:Time | 0.29 | 2 | 48.79 | 0.75 |
| Strain:ROI | 0.73 | 1 | 48.04 | 0.4 |
| ***Time:ROI*** | ***17.19*** | ***2*** | ***48.04*** | ***< 0.001*** |
| Strain:Time:ROI | 1.21 | 2 | 48.04 | 0.31 |

MTT

| Term | F | Df | Residual df | *p* |
|---|---|---|---|---|
| Strain | 0.73 | 1 | 9.94 | 0.41 |
| ***Time*** | ***78.75*** | ***2*** | ***48.59*** | ***< 0.001*** |
| ***ROI*** | ***28.02*** | ***1*** | ***48.03*** | ***< 0.001*** |
| ***Strain:Time*** | ***4.23*** | ***2*** | ***48.68*** | ***0.02*** |
| ***Strain:ROI*** | ***8.06*** | ***1*** | ***48.03*** | ***0.007*** |
| Time:ROI | 2.55 | 2 | 48.03 | 0.089 |
| Strain:Time:ROI | 0.26 | 2 | 48.03 | 0.77 |

**Supplementary Table II - Post-hoc analyses of cerebral perfusion indices before and after tPA**

CBF

| Strain | Time | contrast | Effect size | Estimate | SE | Resid df | t | *p* |
|---|---|---|---|---|---|---|---|---|
| ***SHR*** | ***Occlusion*** | ***Perilesional - Core*** | ***2.09*** | ***45.07*** | ***12.45*** | ***48.02*** | ***3.62*** | ***0.004*** |
| NR | Occlusion | Perilesional - Core | 1.03 | 22.28 | 13.63 | 48.02 | 1.63 | 0.65 |
| ***SHR*** | ***Day 1*** | ***Perilesional - Core*** | ***2.07*** | ***44.67*** | ***12.45*** | ***48.02*** | ***3.59*** | ***0.005*** |
| ***NR*** | ***Day 1*** | ***Perilesional - Core*** | ***1.64*** | ***35.4*** | ***12.45*** | ***48.02*** | ***2.84*** | ***0.039*** |
| SHR | Day 7 | Perilesional - Core | -1.25 | -26.93 | 12.45 | 48.02 | -2.16 | 0.21 |
| NR | Day 7 | Perilesional - Core | -0.6 | -12.96 | 12.45 | 48.02 | -1.04 | > 0.99 |

CBV

| Strain | Time | contrast | Effect size | Estimate | SE | Resid df | t | *p* |
|---|---|---|---|---|---|---|---|---|
| ***SHR*** | ***Occlusion*** | ***Perilesional - Core*** | ***2.02*** | ***0.44*** | ***0.13*** | ***48.04*** | ***3.49*** | ***0.006*** |
| NR | Occlusion | Perilesional - Core | 1.41 | 0.31 | 0.14 | 48.04 | 2.24 | 0.18 |
| SHR | Day 1 | Perilesional - Core | 1.21 | 0.27 | 0.13 | 48.04 | 2.1 | 0.25 |
| ***NR*** | ***Day 1*** | ***Perilesional - Core*** | ***1.73*** | ***0.38*** | ***0.13*** | ***48.04*** | ***2.99*** | ***0.026*** |
| ***SHR*** | ***Day 7*** | ***Perilesional - Core*** | ***-1.95*** | ***-0.43*** | ***0.13*** | ***48.04*** | ***-3.37*** | ***0.009*** |
| NR | Day 7 | Perilesional - Core | -0.72 | -0.16 | 0.13 | 48.04 | -1.25 | > 0.99 |

MTT (contrast = Strain)

| Time | ROI | contrast | Effect size | Estimate | SE | Resid df | t | *p* |
|---|---|---|---|---|---|---|---|---|
| Occlusion | Core | SHR - NR | 0.02 | 0 | 0.11 | 51.98 | 0.03 | > 0.99 |
| ***Day 1*** | ***Core*** | ***SHR - NR*** | ***1.97*** | ***0.31*** | ***0.1*** | ***50.38*** | ***3.1*** | ***0.019*** |
| Day 7 | Core | SHR - NR | 0.85 | 0.14 | 0.1 | 50.38 | 1.34 | > 0.99 |
| Occlusion | Perilesional | SHR - NR | -1.32 | -0.21 | 0.11 | 51.98 | -1.98 | 0.32 |
| Day 1 | Perilesional | SHR - NR | 0.19 | 0.03 | 0.1 | 50.38 | 0.3 | > 0.99 |
| Day 7 | Perilesional | SHR - NR | -0.1 | -0.02 | 0.1 | 50.38 | -0.15 | > 0.99 |

MTT (contrast = ROI)

| Strain | Time | contrast | Effect size | Estimate | SE | Resid df | t | *p* |
|---|---|---|---|---|---|---|---|---|
| ***SHR*** | ***Occlusion*** | ***Perilesional - Core*** | ***-2.71*** | ***-0.43*** | ***0.09*** | ***48.03*** | ***-4.7*** | ***< 0.001*** |
| NR | Occlusion | Perilesional - Core | -1.37 | -0.22 | 0.1 | 48.03 | -2.17 | 0.21 |
| ***SHR*** | ***Day 1*** | ***Perilesional - Core*** | ***-1.81*** | ***-0.29*** | ***0.09*** | ***48.03*** | ***-3.13*** | ***0.018*** |
| NR | Day 1 | Perilesional - Core | -0.02 | 0 | 0.09 | 48.03 | -0.04 | > 0.99 |
| SHR | Day 7 | Perilesional - Core | -1.32 | -0.21 | 0.09 | 48.03 | -2.29 | 0.16 |
| NR | Day 7 | Perilesional - Core | -0.37 | -0.06 | 0.09 | 48.03 | -0.64 | > 0.99 |

**Supplementary Table III - Interaction analyses - consecutive comparisons of contrasts before and after tPA**

CBF

| Time contrast | ROI contrast | Strain | Effect size | Estimate | SE | Resid df | t | *p* |
|---|---|---|---|---|---|---|---|---|
| Day 1 - Occlusion | Perilesional - Core | SHR | -0.02 | -0.4 | 17.6 | 48.02 | -0.02 | > 0.99 |
| ***Day 7 - Day 1*** | ***Perilesional - Core*** | ***SHR*** | ***-3.32*** | ***-71.6*** | ***17.6*** | ***48.02*** | ***-4.07*** | ***< 0.001*** |
| Day 1 - Occlusion | Perilesional - Core | NR | 0.61 | 13.13 | 18.46 | 48.02 | 0.71 | 0.96 |
| ***Day 7 - Day 1*** | ***Perilesional - Core*** | ***NR*** | ***-2.24*** | ***-48.36*** | ***17.6*** | ***48.02*** | ***-2.75*** | ***0.017*** |

CBV

| Time contrast | ROI contrast | Strain | Effect size | Estimate | SE | Resid df | t | *p* |
|---|---|---|---|---|---|---|---|---|
| Day 1 - Occlusion | Perilesional - Core | SHR | -0.8 | -0.18 | 0.18 | 48.04 | -0.99 | 0.66 |
| ***Day 7 - Day 1*** | ***Perilesional - Core*** | ***SHR*** | ***-3.16*** | ***-0.69*** | ***0.18*** | ***48.04*** | ***-3.87*** | ***< 0.001*** |
| Day 1 - Occlusion | Perilesional - Core | NR | 0.31 | 0.07 | 0.19 | 48.04 | 0.36 | > 0.99 |
| ***Day 7 - Day 1*** | ***Perilesional - Core*** | ***NR*** | ***-2.45*** | ***-0.54*** | ***0.18*** | ***48.04*** | ***-3*** | ***0.009*** |

MTT

| Strain contrast | ROI contrast | Time | Effect size | Estimate | SE | Resid df | t | *p* |
|---|---|---|---|---|---|---|---|---|
| NR - SHR | Perilesional - Core | Occlusion | 1.34 | 0.21 | 0.14 | 48.03 | 1.57 | 0.12 |
| ***NR - SHR*** | ***Perilesional - Core*** | ***Day 1*** | ***1.78*** | ***0.28*** | ***0.13*** | ***48.03*** | ***2.19*** | ***0.034*** |
| NR - SHR | Perilesional - Core | Day 7 | 0.95 | 0.15 | 0.13 | 48.03 | 1.16 | 0.25 |

**Supplemental table IV: fractions of hypo- and hyperperfusion regressed against sensorimotor deficit score**

| | CBF | | CBV | | MTT | |
|---|---|---|---|---|---|---|
| Hypoperfusion (%) | 0.0039 | | -0.0028 | | **0.0177*** | |
| | (0.64) | | (0.81) | | **(0.04)** | |
| Hyperperfusion (%) | | -0.0053 | | 0.012 | | -0.028# |
| | | (0.60) | | (0.43) | | (0.06) |
| Strain (hypertensive) | -0.59 | -0.57 | -0.66 | -0.75 | -0.66 | -0.42 |
| | (0.29) | (0.31) | (0.25) | (0.20) | (0.27) | (0.48) |
| Initial ischemic volume | 0.0052 | 0.0050 | 0.0066# | 0.0080* | 0.0041 | 0.0042 |
| | (0.12) | (0.14) | (0.06) | (0.03) | (0.18) | (0.15) |
| AIC | 49.8 | 49.7 | 49.9 | 49.3 | 45.7 | 45.7 |
| RMSE | 1.39 | 1.38 | 1.42 | 1.38 | 1.12 | 1.03 |
| Nagelkerke's R2 | 0.5 | 0.5 | 0.49 | 0.53 | 0.74 | 0.74 |
| # p < 0.1, * p < 0.05, ** p < 0.01, *** p < 0.001 | | | | | | |

*Note: AIC = Akaike Information Criterion; RMSE = Root Mean Square Error; CBF = Cerebral Blood Flow; CBV = Cerebral Blood Volume; MTT = Mean Transit Time*
